# Gradients of Rac1 nanoclusters support spatial patterns of Rac1 signaling

**DOI:** 10.1101/131227

**Authors:** Amanda Remorino, Simon De Beco, Fanny Cayrac, Fahima Di Federico, Gaetan Cornilleau, Alexis Gautreau, Maria Carla Parrini, Jean-Baptiste Masson, Maxime Dahan, Mathieu Coppey

## Abstract

The dynamics of the cytoskeleton and cell shape relies on the coordinated activation of RhoGTPase molecular switches. Among them, Rac1 participates to the orchestration in space and time of actin branching and protrusion/retraction cycles of the lamellipodia at the cell front during mesenchymal migration. Biosensor imaging has revealed a graded concentration of active GTP-loaded Rac1 in protruding regions of the cell. Here, using single molecule imaging and super-resolution microscopy, we reveal an additional supramolecular organization of Rac1. We find that, similarly to H-Ras, Rac1 partitions and is immobilized into nanoclusters of 50-100 molecules each. These nanoclusters assemble due to the interaction of the polybasic tail of Rac1 with the phosphoinositide lipids PIP2 and PIP3. The additional interactions with GEFs, GAPs, downstream effectors, and possibly other partners are responsible for an enrichment of Rac1 nanoclusters in protruding regions of the cell. Using optogenetics and micropatterning tools, we find that activation of Rac1 leads to its immobilization in nanoclusters and that the local level of Rac1 activity matches the local density of nanoclusters. Altogether, our results show that subcellular patterns of Rac1 activity are supported by gradients of signaling nanodomains of heterogeneous molecular composition, which presumably act as discrete signaling platforms. This finding implies that graded distributions of nanoclusters might encode spatial information.

**Significance statement:** The plasma membrane of eukaryotic cells is a highly organized surface where hundreds of incoming signals are transduced to the intracellular space. How cells encode faithfully this myriad of signals is a fundamental question. Here we show that Rac1, a critical membrane-bound protein involved in the regulation of cytoskeletal dynamics, forms small aggregates together with other regulating proteins. These supramolecular assemblies, called nanoclusters, are the “quantal” units of signaling. By increasing the local concentration, nanoclusters set thresholds for downstream signaling and ensure the fidelity of information transduction. We found that Rac1 nanoclusters are distributed as spatial gradients matching the patterns of Rac1 activity. We propose that cells can encode positional information through distributed signaling quanta, hereby ensuring spatial fidelity.

## Introduction

Cell migration and tissue invasion have important roles in cancer metastasis and embryonic development. Among the different mechanisms of migration, protrusion-based mesenchymal migration involves the formation of structures called lamellipodia that alternate between protruding and retracting cycles through actin polymerization and depolymerization (*1*). The regulation of this highly dynamic and adaptable mechanism of motion dictates the outcome of many cellular processes. For example, the stiffness of the branched actin network (*2*), the frequency of its oscillations (*3*), the relative ratio of elongation and branching (*4*) and membrane trafficking (*5*) can be tuned to yield distinct phenotypic effects. This regulation is achieved through a complex coordination of many signaling pathways in which RhoGTPases, small molecular switches that integrate multiple inputs to orchestrate the dynamics of the cytoskeleton, play a pivotal role.

One of the most studied RhoGTPases, Rac1, is at the core of signaling pathways regulating cell polarization and migration. Rac1 is activated and deactivated at the plasma membrane, and possibly at endomembranes, through the interaction with Guanine nucleotide exchange factors (GEFs) and GTPase-activating protein (GAPs) respectively. Rac1 shuttles to and from the plasma membrane through its interaction with Rho GDP-dissociation inhibitors (GDIs), which mask its prenyl group. Rac1 presents spatiotemporal patterns of activity (*6*)(*7*) which extend over few micrometers and last for few minutes during cell migration (*8*). Localized shuttling of Rac1 by GDIs and localized activation by GEFs are two mechanisms capable of producing and maintaining activation profiles. They represent different layers of regulation and their relative importance is still not clear (*9*)(*10*).

Modeling studies (*11*) have identified three main variables controlling the spatiotemporal properties of its subcellular gradients of activation: the spatial distribution of activators and deactivators (GEFs and GAPs, respectively), the cycling rates between activation states and the diffusivity of RhoGTPases at the membrane. Assuming a sharply localized GEF and a uniform GAP distribution, the spatial extent of active Rac1 simply depends on its lifetime in the GTP-bound state and its lateral diffusion coefficient. Yet, we do not know whether the spatial extent of Rac1 activity gradients in the cell, generated by a specific distribution of activators and deactivators, are maintained due to low mobility or short lifetimes. Some of the mechanisms that localize GEFs and GAPs have been identified and described (reviewed in (*8*)). Lipid-interaction domains with varying lipid specificity, BAR domains, tyrosine kinases, scaffold proteins, adhesion complexes and the cytoskeleton have been shown to selectively direct GEFs and GAPs to different plasma membrane (PM) subdomains. In contrast, only few works have focused on the study of RhoGTPases diffusivities (*12*)(*13*)(*14*) or on the determination of cycling rates (*15*)(*16*).

In addition to the molecular parameters encoding the cellular-scale patterns of Rac1 activity, there might be a supramolecular organization of Rac1 signaling not accessible by conventional microscopy. In the last decade, several studies have reported the existence of nanoclusters for membrane-bound signaling proteins (*28*). It has been argued that all signaling proteins might be regulated through nanoclusters (*29*). These nanoclusters accumulate around 10 proteins in less than 250 nm^2^ areas (*17*), producing highly localized increase of concentrations that allow putative thresholds to be overcome. As such, their assumed function is to ensure the transduction of signals with high fidelity, each nanocluster acting as discrete signal processing units digitalizing the input (*18*). The small G-protein Ras presents the best studied case of nanoclustering (*17*). On the plasma membrane, ca. 44% of Ras proteins are organized into ∼9nm nanoclusters composed of 4-7 proteins and having a 0.1-1 s lifetime (*19*). Active and inactive forms of Ras are segregated into different nanoclusters. Ras proteins exist in different isoforms, the H-Ras, N-Ras and K-Ras. They differ in their lipid anchor and yield nanoclusters of varying acidic phospholipid, cholesterol and scaffold proteins composition. As a consequence, they behave differently when the plasma membrane is perturbed through cholesterol depletion or cytoskeleton disruptions (*20*), highlighting the importance of polybasic sequences in proper signal propagation (*21*). In addition, positively charged membrane anchored proteins have been shown to induce PIP2 nanoclustering by charge stabilization (*22*) and equivalent effects for PIP3 have been proposed (*23*). PIP3 and PIP2 are important signaling molecules (*32*), which define the spatiotemporal characteristics of the actin polymerization response (*1*). Similarly, the membrane interacting domain of Rac1 is built up of an unspecific geranylgeranyl isoprenoid lipid and a repetition of basic residues that confer specificity for the negatively charged lipids PIP2, PIP3 (*24*) and phosphatidylserine (*25*)(*26*)(*27*). Yet, despite its fundamental role, it is still unknown whether the RhoGTPase Rac1 forms nanoclusters.

In this work, we used single molecule localization microscopy in live cells (SPT-PALM) (*30*) to address the architecture and dynamics of Rac1 in the basal plasma membrane of NIH 3T3 cells. We found that Rac1 displays static and diffusing states and that Rac1 immobilization is mainly due to its partitioning into nanoclusters. Rac1 immobilization and nanoclustering are enhanced at the front of the cell and correlate with regions of high Rac1 activity. The polybasic anchor of Rac1 is sufficient to drive nanocluster formation but results obtained from Rac1 mutants show that interactions with GEFs, GAPs and effectors are required to enrich nanoclusters at the front of the cell. Using optogenetics combined with single molecule imaging, we causally established that activation of cycling wild-type Rac1 leads to its immobilization and that interactions with effectors are the most efficient in promoting Rac1 immobilization, similarly to what have been observed with H-Ras (*31*). Two-color super-resolution images confirmed that nanoclusters at the active front of the cell are composed of at least Rac1, PIP3 and the WAVE nucleation promoting factor. Additionally, by quantitatively comparing the profiles of Rac1 activity and immobilization in micro-patterned cells, we found that the fraction of Rac1 immobilization is a non-linear function of its activity supporting the existence of an amplification mechanism by which active Rac1 is further immobilized in regions of high Rac1 activity. We propose that interactions with downstream effectors such as WAVE are responsible for this amplification by stabilizing nanoclusters and thus enhancing their lifetime.

Altogether, the heterogeneous composition of Rac1 nanoclusters suggest that they operate as signaling platforms where GEFs, GAPs and effectors are concentrated and where the size of nanoclusters is tightly regulated by cycling between active and inactive states. Importantly, our results show for the first time the existence of subcellular gradients of nanocluster. Their distribution matches the activity measured by a FRET biosensor, suggesting that a supramolecular level of organization mediates Rac1 signal transduction.

## Results

### Rac1 forms nanoclusters

Single molecule tracking experiments have been used in the past to study the diffusivity of Rac1 in spreading MCF7 cells (*14*), dendritic spines (*13*) and within focal adhesion points of Hela cells (*12*). In the present work, we used a single particle tracking photoactivated localization microscopy (SPT-PALM) (*30*) approach in a total internal reflection microscopy (TIRF) configuration. TIRF microscopy allowed us to capture only molecules present in the basal membrane without the noisy contribution of cytoplasmic proteins. The benefit of SPT-PALM approaches is to yield individual localizations in live cells that can be used both to access the supramolecular organization of molecules in the membrane and to build individual trajectories revealing the mobility of the tagged-proteins, as schemed in **figure 1A**. Here, we used live cells stably expressing Rac1 labeled with the photoconvertible protein mEOS2, which we sparsely photoactivated to image single molecules (**figure 1B** and **supplementary movie 1**). Localizations and trajectories were used to build quantitative reporters of the architecture and dynamics of Rac1 at the basal plasma membrane (**figure 1C**).

**Figure 1:**
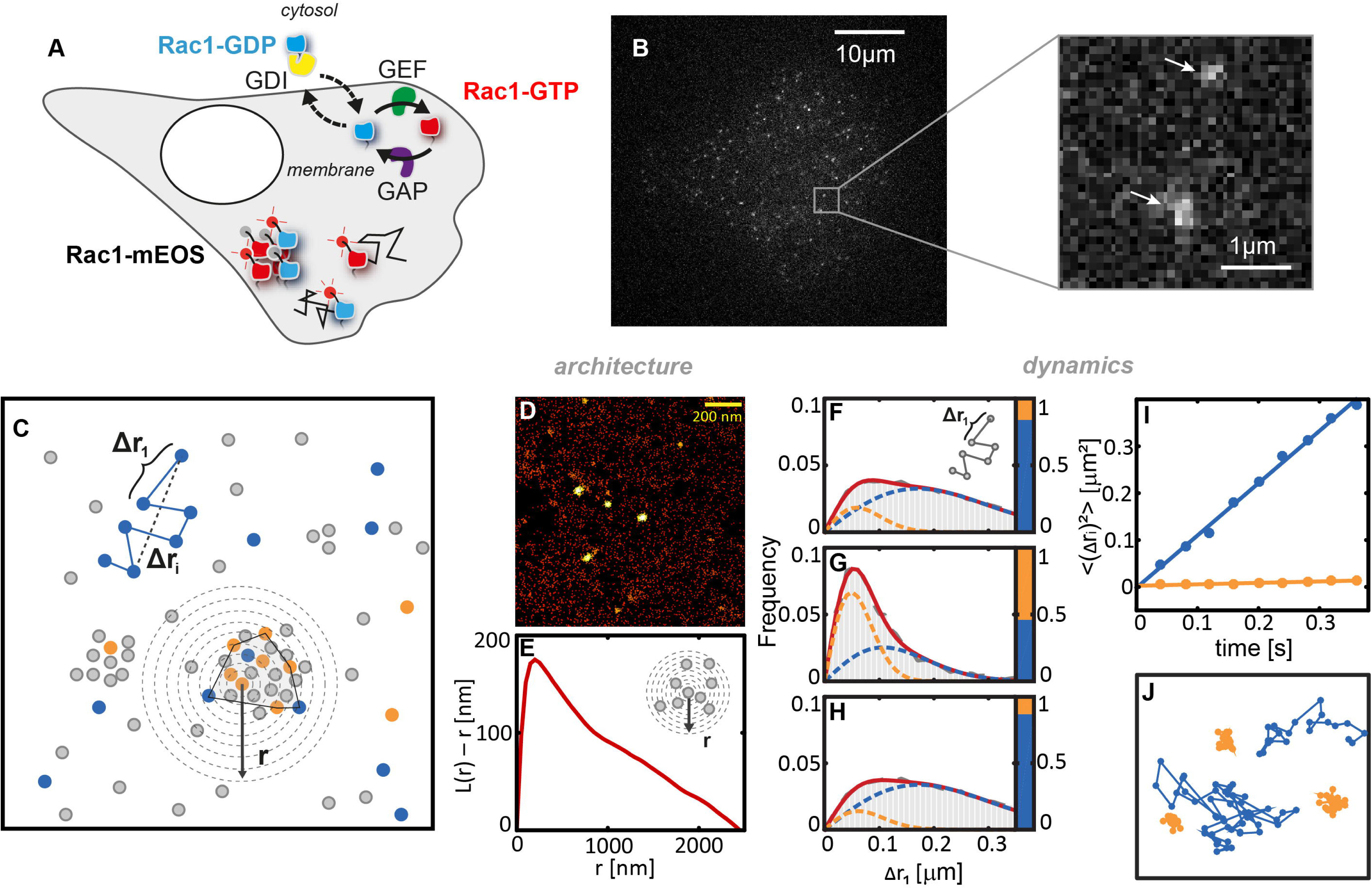
Rac1 forms nanoclusters and presents two diffusive states. **(A)** Scheme of the mechanisms regulating Rac1 activity inside the cell. Rac1 switches between GDP (blue) and GTP (red) loaded forms and shuttles between the membrane and the cytosol. We used a TIRF SPT-PALM strategy, by fusing the photoactivable mEOS2 fluorescent protein to Rac1. Using low power of activation, only few mEOS2 molecules are photoconverted giving access to localizations and trajectories of single Rac1 molecules (either GTP or GDP loaded). **(B)** Example of a frame from a single movie of mEOS2-Rac1-WT at the basal plasma membrane (see **supplementary movie 1**) and zoom showing two individual molecules (arrows). **(C)** Scheme of the parameters extracted from the single molecule movies. Blue/orange (diffusing/immobile) spots are mEOS2 molecules that are imaged and localized from the movies. Gray spots represents mEOS2 molecule that are not imaged. From the trajectories, we extracted Δ*r*_*i*_ the displacement for a time lag *t*_*i*_. From the localizations, we calculated the local density as a function of ***r***, the distance from the center of a molecule (Ripley function). We identified nanoclusters using a DBSCAN algorithm. Nanoclusters were segmented using the convex hull (dashed polygon). A **(D)** PALM image (colorbar 0-60 neighbors) of Rac1 reveals its nanocluster organization that yields a peak in the **(E)** Ripley function. **(F-H)** Single-translocation histograms (grey), Δ*r*_1_, between consecutive frames in the **(F)** whole cell, **(G)** inside nanoclusters and **(H)** outside nanoclusters cannot be fitted with a single Brownian population. When fitted with two states they yield a (orange) quasi-static component and a (blue) freely moving one with different population sizes. The sum of the two components is given by the red curve. Inside nanoclusters, the amount of immobilization represented by the bar graph on the right side of plots in is much higher than outside. **(I)** The mean square displacement recovered from histograms fits are linear with increasing time interval **(figure S4**) and their slopes yield diffusion coefficients of D_mobile_ = 0.28 μm^2^/s for the mobile state and D_static_ = 0.008 μm^2^ /s for the static state. The origin of the mobile state line yields a localization precision 31±3 nm. **(J)** Representative trajectories of the two populations. Details on the methods can be found in the methods section.

A density-based representation of PALM live cell images of wild type Rac1 tagged with mEOS2 (mEOS2-Rac1-WT) revealed that Rac1 forms nanoclusters (**figure 1D**), similarly to Ras proteins(*33*)(*34*)(*20*). Analysis of the spatial distribution of mEOS2-Rac1-WT using a Ripley K-function (L(r)-r) (*35*) (**figure 1E**) and a PC-PALM approach (*36*) provided quantitative supports of nanocluster formation. Ripley functions (**figure 1E**) exhibit a peak at 200nm indicating an inhomogeneous distribution of proteins on the membrane with structures of length scales in the order of hundreds of nanometers. Moreover, fitting of the pair correlation function of mEOS2-Rac1-WT (**Figure S1**) required two components: a Gaussian one corresponding to the localization accuracy associated with multiple observations of the same molecule and an exponential one decaying over a length scale of 100-200 nm, which accounts for the existence of nanoclusters. In contrast, the pair correlation function (**Figure S1**) of a transmembrane domain control (*37*) tagged with mEOS2 can be properly fitted with only the Gaussian component. To further exclude the eventuality of spurious nanocluster identification due to the consecutive imaging of the same immobile protein, we corrected PALM images of mEOS2-Rac1-WT on fixed cells by eliminating the localizations that were within the localization precision in consecutive frames. The corrected images yielded virtually identical images than the uncorrected ones (**Figure S1**).

We next checked if Rac1 nanoclusters are also present for endogenous Rac1. We acquired “Stochastic optical reconstruction microscopy” (STORM) images in fixed cells of immunolabeled endogenous Rac1. Resulting images also show nanoclusters (**figure S2**) and show pair correlation functions that cannot be fitted solely with a Gaussian component (**Figure S1**). Altogether, our measurements provide strong evidence that Rac1 forms nanoclusters at the plasma membrane.

### Rac1 is immobilized in nanoclusters

As previously shown for Ras, nanoclusters can arrest proteins and thus modulate lateral diffusivity on the membrane. We thus assessed if nanoclusters also immobilized Rac1 molecules. We extracted trajectories of single molecules of mEOS2-Rac1-WT (see methods) and built histograms of the displacements of molecules between consecutive frames (**figure 1F-H**), called single translocations hereafter. Such histograms, could not be fitted (*38*) with a model of a single Brownian species and required two populations. The analysis of the distribution of single displacements for increasing time intervals clearly supported the bimodality of the diffusion (**figure 1F-H** and **S3**). The diffusivity of the slower state (*D*_*slow*_ = 0.008 ± 0.003 μm^2^/s) is within the localization precision of our experimental system and can be considered as static. In the rapid state, the diffusion coefficient is *D*_*fast*_ = 0.28 ± 0.003 μm^2^/s, in agreement with the lateral diffusion coefficient of a freely moving membrane-bound protein. Trajectories shown in **figure 1J** are representative of each state of Rac1 mobility.

We looked for a preferential partitioning of the static state in nanoclusters. Nanoclusters were identified and segmented using a density based scanning algorithm (*39*) such that trajectories could be sorted as belonging or not to nanoclusters (see methods). Histograms of single translocations in **figure 1G** show that trajectories within nanoclusters present a fivefold higher static population than those in regions outside nanoclusters (**figure 1H**). We estimated that 15% of all mEOS2-Rac1-WT immobilizations happen inside nanoclusters (**figure S4**). While this number might be largely underestimated because nanoclusters of smaller sizes are missed by our method, this result shows that partitioning into nanoclusters is one mechanism by which Rac1 becomes immobilized.

### Active Rac1 presents an increased fraction of immobilization and nanoclustering

We next assessed the relationship between activation and immobilization of Rac1 by examining the diffusivity and nanocluster partitioning of different Rac1 mutants. **Figure 2A** shows single translocation histograms of mEOS2 tagged wild type Rac1 (mEOS2-Rac1-WT), constitutively active Rac1 (mEOS2-Rac1^Q61L^), dominant negative Rac1 (mEOS2-Rac1^T17N^), and the CAAX-polybasic region that works as a membrane anchor after post-translational modifications. **Figure 2B** shows the distribution of the static populations sizes obtained from fitting the translocation histograms (*38*). Interestingly, the polybasic membrane anchor presents a similar degree of immobilization and nanoclustering as the mEOS2-Rac1-WT, suggesting that nanocluster formation is inherent to the Rac1 CAAX-polybasic C-terminals domain of the protein. This phenomenon, is consistent with previous reports on the capacity of the C-terminal polybasic domain to mediate Rac1 oligomerization (*40*). However, mEOS2-Rac1^Q61L^ which has the largest static population (**figure 2B**), the highest peak in Ripley K-functions (**figure 2C**) and the highest percentage of localizations within clusters (**figure 2D**), shows that immobilization and nanoclustering have a positive correlation with Rac1 activity. These results, together with previous reports (*12*)(*14*)(*13*), provide robust evidence that in migrating fibroblasts GTP-loaded active Rac1 is less mobile than its inactive counterpart.

**Figure 2:**
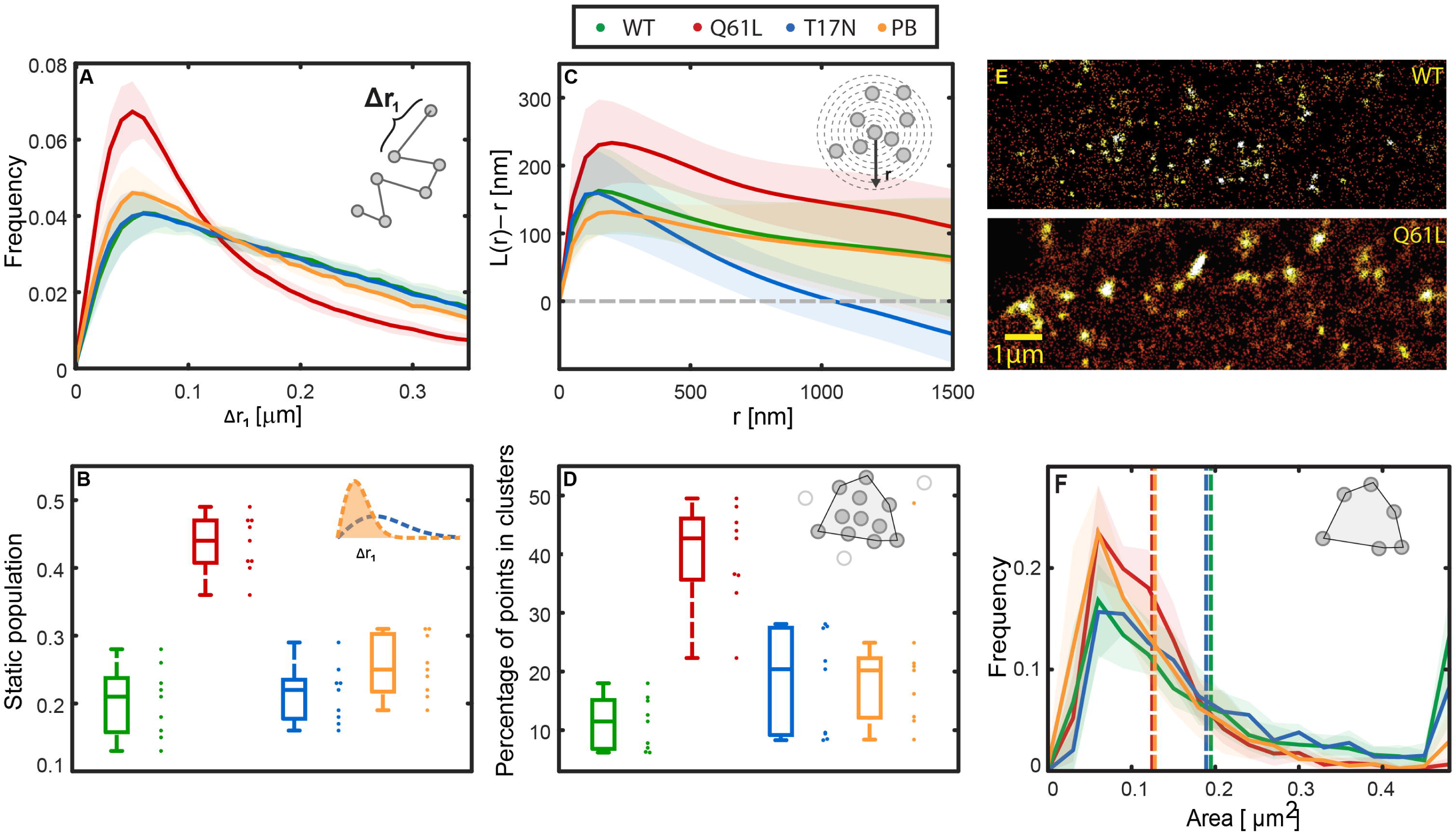
Active Rac1 present a decreased diffusivity and an increased nanoclustering. **(A)** Single translocation histograms obtained from single molecule movies of (mEOS2-Rac1-WT, green) wild type, (mEOS2-Rac1^Q61L^, red) active and (mEOS2-Rac1^T17N^, blue) inactive Rac1 mutants and the (PBCAAX, orange) polybasic CAAX control membrane anchor. The histograms are fitted with two independent populations of different diffusivity (see methods). The **(B)** integrated relative area of the static population obtained in **(A)** is larger for mEOS2-Rac1^Q61L^ than that of mEOS2-Rac1-WT, mEOS2-Rac1^T17N^ and the PBCAAX membrane anchor showing that the degree of immobilization increases with Rac1 activity. The peak in the **(C)** Ripley function L(r)-r, measuring the degree of Rac1 nanoclustering, is higher for mEOS2-Rac1^Q61L^ showing that increasing activity of Rac1 produces, as well, higher nanoclustering. The **(D)** ratio of points contained within nanoclusters obtained with a dbscan algorithm is more than twice larger for mEOS2-Rac1^Q61L^. **(E)** PALM images (colorbar 0-60 neighbors) of representative nanoclusters of (top) mEOS2-Rac1-WT and (bottom) mEOS2-Rac1^Q61L^ exhibit a significant difference in nanocluster sizes. Mean **(F)** nanocluster surface areas shown in dashed lines are larger for mEOS2-Rac1^Q61L^ and mEOS2-Rac1^T17N^. (A) and (F) are means of 9 different single-cell histograms and errobars are calculated as standard deviations. The mean Ripley function in (B) is a mean of 9 different single-cell Ripley functions with errobars calculated as the standard deviation. Boxplots in (B) and (D) represent the median of measurements on 9 different cells.

In addition to the differences in nanocluster partitioning among Rac1 mutants, PALM images **(figure 2E)** of representative nanoclusters for mEOS2-Rac1-WT and mEOS2-Rac1^Q61L^ show a clear difference in size. mEOS2-Rac1-WT displays nanocluster sizes comparable to the ones of the polybasic-CAAX membrane anchor, whereas mEOS2-Rac1^T17N^ and mEOS2-Rac1^Q61L^ display twice larger nanoclusters (**figure 2F**). The quantification of the number of proteins per nanoclusters is a difficult task due to the blinking of mEOS2 (*41*)(*42*)(*42*). However, on average, we estimated that mEOS2-Rac1^Q61L^ and mEOS2-Rac1^T17N^ mutants present 233 ± 110 and 232 ± 49 localizations per nanoclusters, whereas mEOS2-Rac1-WT and the polybasic CAAX anchor present 97 ± 33 and 83 ± 38 localizations. The localizations can be used as a loose estimate of the real number of molecules per nanocluster. If we consider that, in our experimental conditions, a single molecule is counted on average 2.3 times and that the photophysics of mEOS2 allow sampling of only 78% of the molecules (*41*), the number of molecules per nanoclusters can be estimated as 0.55 times the number of localizations per nanoclusters. Hence mEOS2-Rac1-WT nanoclusters are composed of approximately 50 molecules, about 5 times more than the number of Ras molecules in its nanoclusters (*19*). Larger areas and larger numbers of localizations per nanoclusters present in mEOS2-Rac1^Q61L^ and mEOS2-Rac1^T17N^ mutants show that the cycling between active and inactive states is a major factor regulating nanocluster size.

Cycling rates are of high relevance in signaling. Fast cycling of Rac1, but not locking of Rac1 in its GTP-bound form, was shown to transform cells, like the oncogenic activation of upstream GEFs (*43*) (*16*). These results suggest that the cycling kinetics of Rac1 activation determine the transduction efficiency of Rac1 downstream signaling. Given that mEOS2-Rac1^Q61L^ is locked in its GTP-bound state, we examined whether the link between diffusivity and activity identified in mEOS2-Rac1^Q61L^ was also present in cycling Rac1. To this end, we coupled single molecule tracking experiments with optogenetic activation (*44*) (**figure 3A**). In transiently transfected cos7 cells, we illuminated for 10 minutes a specific region of the cell to recruit at the plasma membrane the catalytic domain of the Rac1 GEF Tiam1 (Cry2-Tiam1-iRFP), thereby inducing localized activation of Rac1. Our optogenetic activations led to a 1.2 to 2.2 fold increase of Cry2-Tiam1-iRFP inside the region of activation (**figure S5**). When analyzing single translocation histograms we found that only mEOS2-Rac1-WT displayed an increase in the static population size upon recruitment of Cry2-Tiam1-iRFP (**figure 3B**). We acquired single molecule movies of all mEOS2 tagged Rac1 mutants and the polybasic-CAAX anchor before and after recruitment (**figure 3C-H**) and we mapped the diffusivities over cells using a recently developed methodology (see methods). The polybasic-CAAX membrane anchor and mEOS2-Rac1^T17N^ cannot engage effectors and mEOS2-Rac1^Q61L^ is already in the active state and cannot increase its interaction with effectors. Taking into account that Cry2-Tiam1-iRFP has a diffusivity of 0.1 ± 0.03 μm^2^/s (*45*) comparable to that of the mobile population of Rac1, we attributed the increased immobilization of mEOS2-Rac1-WT to an increase in the amount of active molecules and the consequent interaction with effectors. These optogenetic experiments show that for cycling Rac1 there is a causal relationship between activation and immobilization.

**Figure 3:**
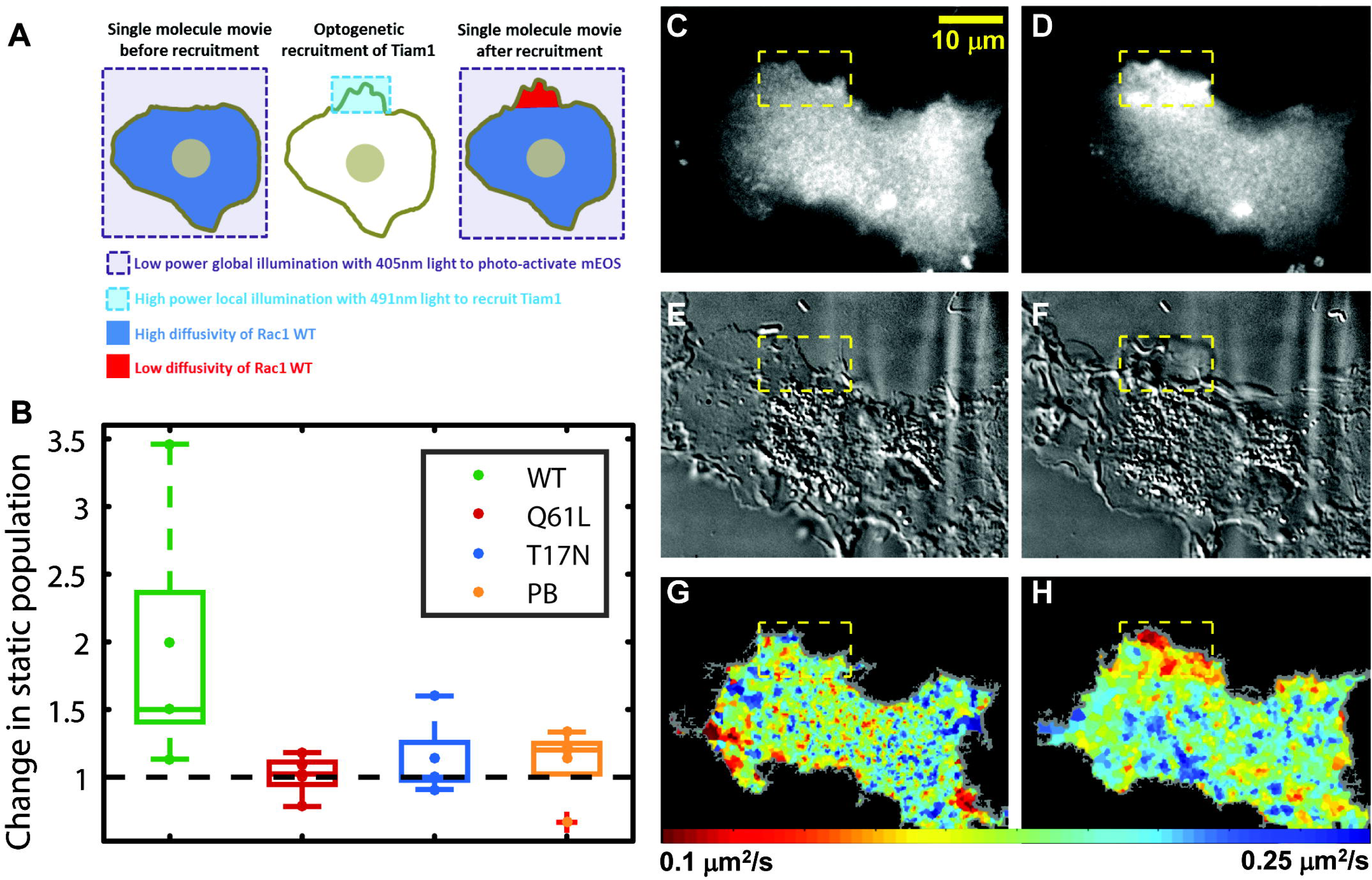
Diffusivities as a function of Rac1 activity modulated through optogenetics. **(A)** Schematic of the experiment. (left) A single molecule movie is acquired by photoconverting mEOS2-Rac1 with low global 405nm illumination to avoid significant optogenetic recruitment. (middle) A 10 min local recruitment step is performed, higher power 491 nm light is used to illuminate a region of interest and recruit Tiam1, a GEF of Rac1, with local specificity. (left) Another single molecule movie is acquired. **(B)** Initial and final single molecule movies were localized and tracked to yield single translocation histograms as the ones shown in figure 1. The ratio of the static population within the activation region between the final and initial movie shows an increase of the immobilization upon optogenetic activation only for mEOS2-Rac1-WT. iRFP channel images **(C)** before and **(D)** after optogenetic activation show Tiam1 recruitment efficiency and DIC images **(E)** before and **(F)** after optogenetic activation expose ruffling induced by Tiam1 recruitment. Diffusivity maps **(G)** before and **(H)** after optogenetic activation exhibit immobilization of mEOS2-Rac1-WT confined to the activation region. 7 cells were used for each condition.

### Rac1 presents similar gradients of immobilization, nanocluster density, and activity

Motivated by previous studies that identified spatial profiles of RhoGTPases activity across cells (*46*), we aimed at comparing them with immobilization profiles and nanocluster distribution. To this end, we plated cells on crossbow fibronectin micropatterns to obtain a normalized cell shape and organization (*47*). The “front” of these cells exhibits ruffling (*48*) and mimics a lamellipodium rich in branched actin. This approach allows the comparison of several measurements taken in different experiments and offers a template for a multiplex mapping approach (**figure 4A)**. Due to the reduced cell-to-cell variability, we were able to average and map in the same referential the fraction of immobile molecules, the nanocluster densities, and the FRET ratiometric images (**figure 4B-E**).

**Figure 4:**
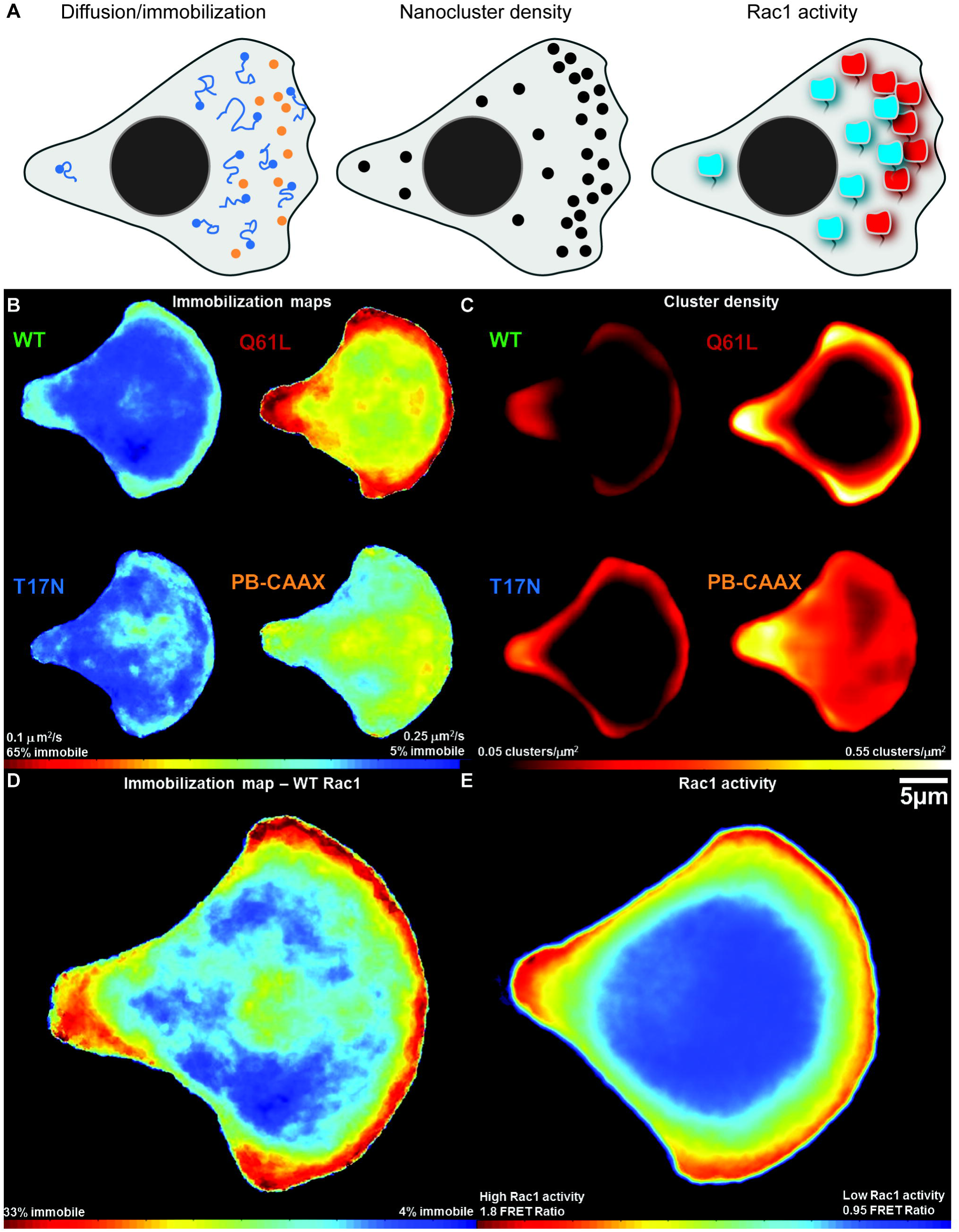
Rac1 diffusivity, activity and nanocluster distribution along normalized polarized cell states imposed by fibronectin crossbow micro-patterns. The cartoon in **(A)** describes the three parameters presented in this figure. **(B)** Immobilization/diffusion maps were obtained from 9 cells per mutant and 18 cells for the mEOS2-Rac1-WT. Cells were tessellated with a voronoi mesh, the local diffusion coefficient in each region was estimated from the single molecule localizations using an inference approach (see methods), and then cells were wrapped onto the average cell shape and averaged (see methods). Immobilization maps of Rac1 mutants show a decreased overall diffusivity for mEOS2-Rac1^Q61L^ and an inhomogeneous diffusivity distribution for all three forms of Rac1 with lower diffusivity at the front and back in contrast to the polybasic anchor that exhibits uniform diffusivities. **(C)** Nanocluster density maps (see methods) show a higher nanocluster density for mEOS2-Rac1^Q61L^ and an increased nanocluster density at the front of the cell for all three Rac1 mutants. Comparison of **(D)** immobilization maps with a maximized dynamic range, **(E)** nanoclusters distribution of wild type Rac1, and **(E)** Rac1 activity maps obtained from FRET biosensor ratios all exhibit a gradient from front to center.

We first acquired single molecules movies (2000-5000 frames at 25 Hz) with densities comparable to **figure 1B** (0.2 molecules/μm^2^). We then mapped Rac1 diffusivity in 9 to 18 individual cells for each mutant and we averaged those maps (**figure 4B**) after morphing each cell onto the average shape (see methods). mEOS2-Rac1-WT, mEOS2-Rac1^Q61L^ and mEOS2-Rac1^T17N^ exhibit diffusivity gradients from the front to the middle of the cell with a region of lowest diffusivity along the cell front-most region. mEOS2-Rac1^Q61L^ presents the greatest contrast in diffusivity between front and middle. Since a given local average diffusion coefficient corresponds to a given local proportion of immobile molecules, diffusivity maps can be interpreted in terms of local fraction of immobilization (see colorbar in **figure 4B,D**). By taking into account the diffusion constant of the slow and fast states derived from tracking experiments, average diffusivities *D*_mean_ = *f*_*i*_. *D*_slow_ + (1 – *f*_*i*_). *D*_fast_ yielded immobilization fractions, *f*_*i*_. In the same single molecule movies, nanoclusters were identified and their spatial densities mapped onto the cell (see methods). As expected given our previous results, the nanocluster density map in **figure 4C** shows that mEOS2-Rac1-WT, mEOS2-Rac1^Q61L^ and mEOS2-Rac1^T17N^ present nanocluster enrichment at the front of the cell, supporting again the link between nanocluster partitioning and immobilization.

We also measured the Rac1 activity map on crossbow micropatterns with a FRET biosensor (*49*). The nanocluster distribution and diffusion map of the WT (**figure 4D,E**) match the biosensor signal (**figure 4F**), all showing a decaying gradient from the front to the center. Thus, in an unperturbed condition, we see a clear positive correlation between Rac1 activity, immobilization and nanocluster density.

### GEF/GAP cycling rather than GDI-mediated membrane shuttling regulates Rac1 activation patterns in spread cells

Localized shuttling of Rac1 to the membrane is one of the processes potentially regulating Rac1 activation. The relative weight of local activation versus local delivery in cell polarity establishment has been addressed before for cdc42 (*9*)(*10*). Localized delivery has been proposed as a the critical mechanism in the establishment of cell polarity in cells minutes after attachment (*14*). Yet, the importance of localized delivery may differ in the context of already spread cells. It was shown that the attachment and spreading processes involve a particular set of signaling pathways (*50*), which may not be triggered once cells have reached a steady state. To evaluate the role of localized delivery in the context of already spread cells, we performed a plasma membrane turnover analysis based on photobleaching experiments.

Fluorescence Recovery After Photobleaching (FRAP) experiments of the green form of mEOS2-Rac1-WT on the whole basal membrane with TIRFM show that the shuttling of Rac1 to the membrane slows down along the spreading process. By performing FRAP experiments 30 minutes and 3 hours after plating, we found that fluorescence recovery times increased from ca. 6 minutes to ca. 20 minutes (**figure S6**). A turnover time of 20 minutes in fully spread cells is of the same order of magnitude as the plasma membrane recycling. This experiment shows that GDI-mediated shuttling occurs on a longer timescale than protrusion/retraction cycles. Therefore, we considered that the predominant mechanism for the generation and maintenance of activation profiles in our experimental conditions was the localized cycling of Rac1 and not its localized delivery.

### Rac1 polybasic tail is sufficient for nanocluster partitioning but interactions with Rac1 partners are required for nanocluster enrichment in active regions of the cell

To further dissect the role of Rac1 molecular interactions in regulating nanocluster distribution, we next quantified the enrichment of nanoclusters in the front of the cell for all mutants, exploiting the fact that they have distinct interacting partners. Among the Rac1 interactome, the best-characterized Rac1 partners are GEFs, GAPs, and the direct effectors. mEOS2-Rac1-WT can interact with all of them. mEOS2-Rac1^Q61L^ can interact with GAPs and effectors and perhaps also with GEFs as demonstrated for Ras proteins (*54*). However, mEOS2-Rac1^T17N^ exhibits high affinity for GEFs but cannot bind effectors or GAPs.

To quantify Rac1 tendency to cluster in different parts of the cell, we divided the crossbow in three different regions as shown in **figure 5A** and measured the density of nanoclusters (**figure 5B**) and the percentage of Rac1 detections in nanoclusters for each region (**figure 5C**). We chose to exclude the back of the cell from the analysis given that its morphology departs from the canonical lamellipodia. Cells plated in crossbow micropatterns present a “small front” at the back, characterized by a high concentration of cortactin (*47*) and high branching. In this aspect, they differ from freely migrating cells that exhibit a retracting tail.

**Figure 5:**
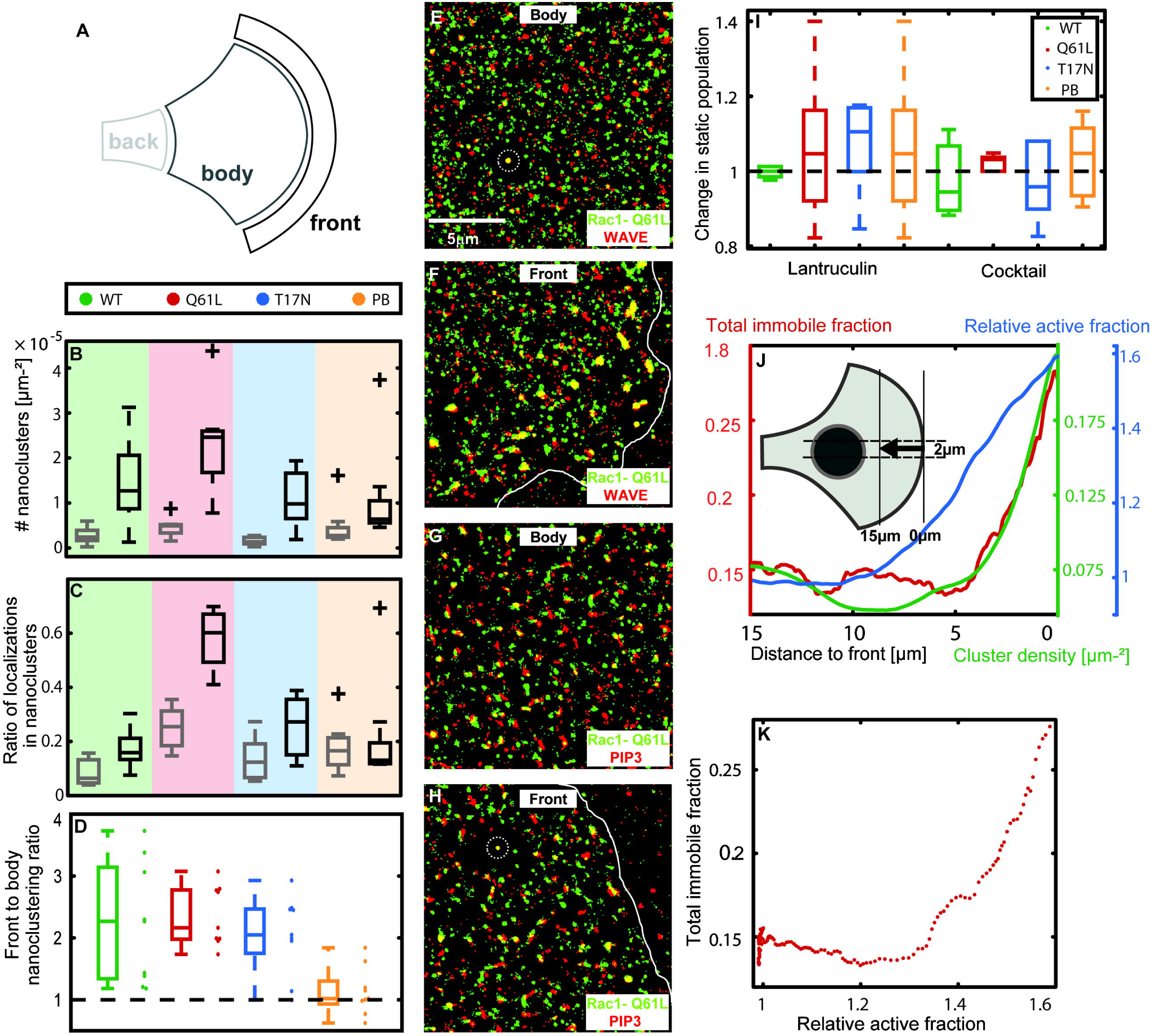
Rac1 nanocluster, activity, immobilization and composition distribution. **(A)** Cells were divided into regions named front, body and back as depicted in the sketch. **(B)** The number of nanoclusters per surface area and **(C)** the fraction of points in nanocluster were measured for each region in each cell and then averaged across 9 cells. Both the number of nanocluster and the total amount of points in nanoclusters is increased at the front of Rac1 mutants in comparison to the body but is comparable for the polybasic CAAX anchor. **(D)** The front to body ratio of points in nanoclusters is ca. 2 for all Rac1 mutants and ca. 1 for the polybasic CAAX anchor. In order to compare immobilization, nanoclustering and activity profiles, we averaged 2μm horizontal stripes across the cell center **(E)** of all three maps. **(E-H)** STORM-PALM images of fixed cells plated in crossbow micropatterns expressing (green) mEOS-Rac1^Q61L^ were constructed using primary antibodies against the **(E,F)** WAVE complex and **(G,H)** PIP3 and (red) secondary antibodies tagged with Alexa Fluo 647 for a front and body regions. The colocalization of mEOS-Rac1^Q61L^ and Alexa 647 is shown in yellow. mEOS-Rac1^Q61L^ and WAVE exhibit nanoclusters of high colocalization at the front of the cell but neglectable colocalization in the body. mEOS-Rac1^Q61L^ also colocalizes with PIP3 but the contrast between front and body is less striking than for WAVE. **(I)** The invariance of the static population fraction for all mutants upon treatment with either lantruculin or a drug cocktail that freezes actin dynamics (*59*) suggest that the formation of Rac1 nanoclusters does not depend on actin. **(J)** Relative Rac1 activity profiles (blue) for the first 15 μm behind the cell edge are twice longer profile than immobile fractions (red) and cluster densities (green) ones. **(K)** Immobilization fractions show a non-linear dependence with relative active fractions.

A similar number of immobilizations as the one reported here has been seen for the polybasic-CAAX motif inside and outside of focal adhesions in Hela (*51*) and MEF (*52*) cells and in dendritic spines (*13*). However, during spreading of MCF7 cells, the polybasic-CAAX anchor does not seem to present a slowly diffusing population (*14*). Here, we identified that 18% of the polybasic-CAAX anchor of Rac1, similarly to that of H-Ras (*53*), is organized into nanoclusters (**figure 5C**) and that 23% of the immobile population can be found within nanoclusters (**figure S4**).

Yet, the interactions responsible for polybasic-CAAX nanocluster formation are insufficient to enrich Rac1 nanoclusters at the front of the cell (**figure 5D**). In contrast, mEOS2-Rac1-WT, mEOS2-Rac1^T17N^ and mEOS2-Rac1^Q61L^ (*54*)(*55*) exhibit a two-fold increase in nanocluster density at the front (**figure 5D**), very likely due to additional interactions. These results suggest that GEFs, GAPs and effectors are sufficient for a relative enrichment of Rac1 nanoclusters at the front of the cell. The significant increase of nanoclustering in mEOS2-Rac1^Q61L^ suggest that interactions with effectors, strongly present in this mutant, seem to be the most effective in promoting nanocluster partitioning.

To test this hypothesis, we looked for the presence within nanoclusters of WAVE2, a major Rac1 effector, and PIP3, which recruits GEFs and GAPs. We acquired two-color PALM/STORM images of cells expressing mEOS2-Rac1^Q61L^ and immunolabeled WAVE2 or PIP3. Supporting our hypothesis, we observed a colocalization of mEOS2-Rac1^Q61L^ and WAVE2 (**Figure 5E-G**) and a colocalization between mEOS2-Rac1^Q61L^ and PIP3 (**Figure 5G-H**) in some of the nanoclusters at the front.

### Rac1 nanoclusters do not depend on the actin cytoskeleton

Actin has been proposed as an inducer of membrane protein nanoclusters either via the formation of transient contractile regions at the plasma membrane that stabilize liquid order domains and couple to extracellular Glycosylphosphatidylinositols-anchored proteins (GPI-Aps), or through the direct interaction of transmembrane proteins with actin filaments (*58*)(*33*). Indeed, nanoclusters of different Ras isoforms exhibit selective dependences on actin. **Figure 5I** shows that treatment with latrunculin and cocktails that freeze actin dynamics (*59*) does not have an effect on the diffusivity of any of the Rac1 mutants or the PB-CAAX anchor control. These results suggest that, like H-RasGTP (*33*)(*60*)(*61*), Rac1 is found in nanoclusters that do not depend on the actin cytoskeleton.

### Partitioning of Rac1 in nanoclusters is amplified in regions of high Rac1 activity

To further assess the role of interactions in nanoclustering, we performed a detailed quantification of Rac1 immobilization fractions, nanocluster density and Rac1 activity at the cell front (**figure 5J**). Immobilization fraction and nanocluster density profiles can be perfectly overlaid, whereas activity gradients (FRET signal) shows a twice-larger spatial extent (**figure 5J**). Plotting the nanocluster density as a function of the activity shows a nonlinear relationship between the two (**figure 5K**). Immobilization fractions are constant for low Rac1 activity. However, for increasing Rac1 activity, the immobilization fraction increases drastically. This observation points to the existence of an amplification mechanism by which active Rac1 molecules have an enhanced propensity to partition into nanoclusters in regions of high Rac1 activity.

## Discussion

We performed a single molecule analysis of Rac1 mobility and supramolecular architecture in migrating fibroblasts. Our main finding is that a significant fraction of Rac1 at the plasma membrane is found in nanoclusters of a few tens of molecules, which are distributed as gradients matching Rac1 subcellular patterns of activity.

Since the polybasic anchor of Rac1 forms nanoclusters and since nanocluster partitioning is independent of actin, Rac1 nanocluster formation is probably driven by electrostatic interactions of its polybasic-CAAX anchor with negatively charged lipids such as PIP2 and PIP3, as previously proposed (*62*). Nanoclusters would form and dissociate spontaneously, without the requirement of active processes or biochemical modifications. Previous studies on the formation of nanoclusters of different Ras isoforms (*20*)(*33*) highlighted the importance of the protein anchor in signaling. These studies identified the role of cholesterol, different membrane anionic lipids, nucleotide load, degree of palmitoylation and protein conformations in the formation and composition of Ras nanoclusters. The anchor of Rac1 resembles K-Ras in the presence of a polybasic region but rather resembles H-Ras in its mono-palmitoylation. Palmitoylation has been shown to induce partitioning of Rac1 into cholesterol-rich liquid-ordered regions (*63*) of sizes in the range of tenths of micrometers. One way to reconcile these data with ours is to consider that nanoclusters belong to larger structures, micrometer-sized, which depend on actin and cholesterol but which do not play a role in Rac1 immobilization.

Supporting the role of charged lipids in Rac1 nanoclustering, PIP2 and PIP3 form nanoclusters in PC12 (*27*)(*64*) and INS-1 (*65*) cells. These lipid nanoclusters are segregated and their diameters are 70nm for PIP2 and 120nm for PIP3. The spatial distribution of PIP3 and PIP2 nanoclusters was not addressed here, but other studies reported non overlapping distributions of PIP3 and PIP2 at the cellular scale (*66*). PIP3 accumulates at the leading edge and adhesions zones during guided cell migration of fibroblasts (*67*) and in membrane protrusions during random cell migration (*68*). In addition, PIP3 directly recruits WAVE to the membrane of polarized cells through a basic sequence in its N-terminal part in an actin independent manner (*56*)(*57*). Our results suggest an additional regulatory function of PIP2 and PIP3, that of inducing nanoclustering of Rac1 via the interaction with its polybasic membrane anchor through coulombic interactions (*62*)(*27*)(*69*).

We found that Rac1 nanoclusters are enriched at the front of the cell, contrarily to the nanoclusters of the polybasic anchor. The subcellular enrichment of nanoclusters is mediated by a second set of interactions, with the GEFs, GAPs, effectors, and possibly other Rac1 partners. In our experiments, the anisotropic spatial cue is given by the asymmetric adhesive crossbow patterns. This constraint yields an organized cell architecture with focal adhesions enriched at the adhesive borders (*47*) that is expected to give rise to an anisotropic distribution of GEFs and GAPs in two different ways. First, direct recruitment and activation of Rac1 to early focal adhesions, the so-called focal complexes at the lamellipodial edge, has been shown to happen via the GEFs β-Pix, DOCK180, Trio, Vav2, Tiam1, and a-Pix (*70*) in a cell type dependent manner. In particular, Tiam1 accumulates at focal complexes of migrating cells and its activation mechanisms has been elucidated (*71*). But also, indirect recruitment and activation of Rac1 in the proximity of focal complexes can happen via PIP3. Indeed, most Rac1 GEFs are recruited with high efficiency by PIP3, but not by other anionic lipids, due to the specificity of PH domains (*72*). The imposed asymmetry in fibronectin, yields an intracellular anisotropy of focal adhesions and a consequent anisotropy of all the signaling components from PIP2 to PIP3, GEFs, GAPs that results in an enrichment of cortactin at the front of crossbow micropatterns (*47*).

Among Rac1 interacting partners, effectors appeared to be the most effective in biasing nanocluster distribution. Indeed, mEOS2-Rac1^Q61L^ presents considerably higher nanoclustering and colocalizes strongly with WAVE in super resolution images. The importance of WAVE in promoting Rac1 nanoclustering can explain the amplification we observed in **figure 5K**. Since the distribution of Rac1 effectors correlates with the local density of nanoclusters, we propose that the enrichment of nanoclusters at the front is due to an increased residence time of active Rac1 within nanoclusters rather than an enhanced seeding of nanoclusters. The amplification mechanism would then operate in the following way: active Rac1 and PIP3 (*56*)(*57*) recruit effectors to nanoclusters that become trapped and are capable of further retaining active Rac1 within nanoclusters. As a result, this mechanism would act as a Rac1 positive feedback. WAVE is also known to concentrate at the very tip of the lamellipodium and would enhance the partitioning of active Rac1 in this region. Consistent with this idea, the accompanying manuscript by Mehidi *et al.* demonstrated that transient Rac1 immobilizations at the lamellipodium tip are correlated with Rac1 activation that triggers membrane protrusion. Nevertheless, the constitutively inactive mEOS2-Rac1^T17N^ and the CAAX-polybasic anchor, which are immobilized in nanoclusters, are not displaying partitioning at the lamellipodium tip, suggesting that the signaling nanocluster do not form at the lamellipodium tip in migrating cells.

In this work, we propose that nanoclusters comprising active Rac1 molecules act as signaling units regulating downstream transduction. Such nanodomains have already been observed for other membrane-bound signaling proteins and several hypotheses have been proposed to explain their functional relevance (*73*). High local concentrations within nanoclusters could set a threshold for signal transduction. Weak interactions can be stabilized by cooperativity in nanoclusters enabling the activation of downstream signaling cascades, as recently shown with the aPKCs kinase transducing intracellular calcium (*28*). For Ras (*74*), it was shown that nanoclusters act as a signal processing step converting analog inputs (concentrations of ligands) into digital ones (numbers of nanoclusters) and giving rise to other analog outputs (levels of intracellular active species) further processed downstream. The functional role of an analog-to digital-to analog processing is not fully understood but it was proposed to provide high fidelity responses (*74*). More recently (*75*), it was proposed that nanoclusters of about 10 molecules exhibit an optimal fidelity. Digitalization reduces the numbers of output states, but also reduces the noise in the system, and a tradeoff between the two maximizes information transmission.

For Rac1, we do not know yet the functional role of nanoclustering, but we can hypothesize that the same concepts hold true. Rac1 nanoclusters may work as a means to generate discrete signals by setting up WAVE thresholds that modulate actin polymerization in a non-linear way, as suggested by the need of coincident anionic lipids, phosphorylation of WAVE and active Rac1 (*76*). In addition, Rac1 nanoclusters may modulate reaction rates by modifying the local concentration of reactants (*77*)(*78*) adding an additional layer of regulation aimed at refining profiles of Rac1 activity and actin polymerization. Along this line of thought, the spatial modulation of cycling rates has been observed in wound healing experiments in oocytes (*79*). Here, even if the spatial distribution of signaling molecules has already been recognized (*80*), we show for the first time that a graded distribution of nanoclusters is a means to provide a spatially modulated digital output.

Nanoclusters can support a double role in generating high fidelity responses. In addition to noise reduction, nanoclusters can help in the maintenance of sharp regions of signaling activity (*81*). Indeed, Rac1 partitioning into nanoclusters is one of the mechanisms through which Rac1 is immobilized and its diffusion spatially restricted. Previous studies (*11*) aimed at characterizing the link between diffusivity, cycling and source distribution showed that decreasing the diffusion constant throughout the cell can enhance the sharpness of activity gradients. Our results show that this effect can be acting through the diffusivity gradients that follow activation profiles from the front to the back of the cell. As seen in **figure S12**, immobilization gradients enable an increase in deactivation time by a factor of ∼2. Even though this might appear a mild increase, we believe that in endogenous conditions the restriction of diffusion might be a significant mechanism to maintain sharp activation gradients since the total fraction of immobile Rac1 might be higher, as suggested by the increased nanoclustering seen for endogenous Rac1 **(figure S7)**.

In conclusion, our findings can be summarized in the model sketched in **figure 6**. Polarized migrating cells exhibit opposite gradients of PIP3/PIP2 with an enrichment of PIP3 at the front and PIP2 in the body (*66*) (*67*)(*68*). Because PIP3 and PIP2 organize in segregated nanoclusters (*27*)(*64*) (*65*), we believe that the front of the cell presents a higher number of PIP3 nanoclusters and the body a higher number of PIP2 ones. The affinity of the polybasic CAAX anchor for either PIP2 or PIP3 might be comparable given that they are based on non-specific coulombic interactions and thus nanoclusters labeled by this anchor or are homogeneously distributed. However, PIP3 nanoclusters at the front recruit GEFs and GAPs and are enhancing the lifetime of Rac1 nanoclusters. Additionally, PIP3 nanoclusters and concomitant WAVE recruitment by GTP-loaded Rac1 (*56*)(*76*) further enhances nanoclusters lifetime and nanocluster enrichment, which would consequently provide a positive feedback mechanism, sustaining cell migration.

**Figure 6:**
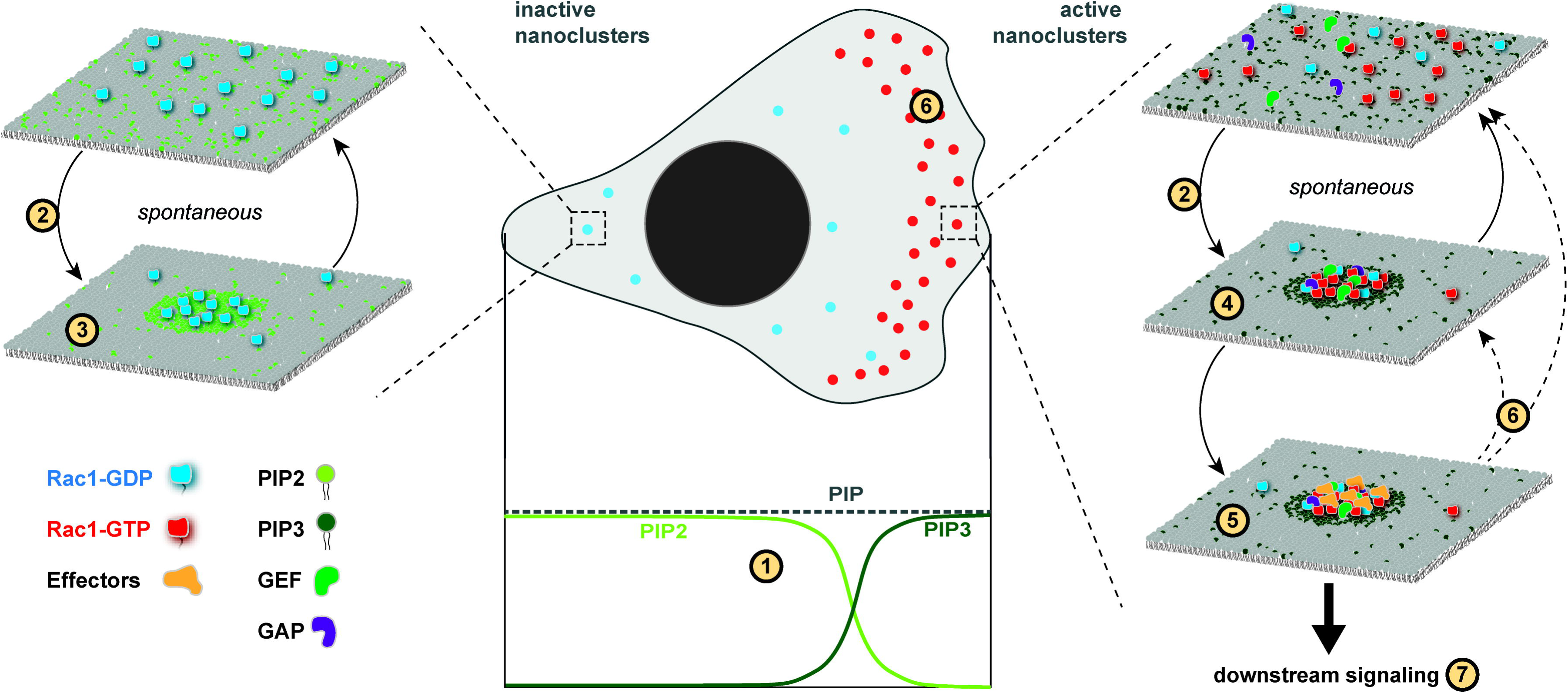
Model for Rac1 nanoclustering. Opposing gradients of PIP3 and PIP2 across the cell (1) and the segregation into different clusters at the molecular level proposes an enrichment of active Rac1-PIP3 nanoclusters at the front. Both active and inactive Rac1 can form nanoclusters spontaneously through electrostatic interactions (2). In the body, inactive Rac1 and PIP2 form inactive nanoclusters (3). At the front, active Rac1 and PIP3 form active nanoclusters (3) which also integrates GEFs and GAPs. These active nanoclusters recruit Rac1 effectors (5) which stabilize nanoclusters lifetime and consequently enrich nanocluster density at the cell front (6). The heterogeneous composition of active nanoclusters suggest the existence of signaling platforms necessary for downstream signaling (7). Under this assumption, the stabilization of nanoclusters by effectors act as a positive feedback to increase the amount of Rac1 signaling where a high density of effectors is present.

## Materials and methods

**Cell culture.** All single molecule tracking, super-resolution experiments, and FRET biosensor imaging were performed on NIH 3T3 cells. Combined single molecule tracking and optogenetics experiments were done with cos-7 cells. In every case, cell culture was performed according to the ATCC proposed protocol, cultured at 37 C in 5% CO_2_ in DMEM (Dulbecco’s modified Eagle’s medium) and supplemented with 10% fetal calf serum. For single molecule tracking and super-resolution experiments we produced lentiviral stable cell lines expressing mEOS2-Rac1 mutants with a pHR backbone plasmid synthesized by Genescript. Cells were sorted using fluorescence-activated cell sorting. Optogenetics experiments were performed via triple transfection of CIBN-GFP (*45*), TIAM_linker_CRY2_IRFP obtained following the same routine as in (*45*), and mEOS2-Rac1 mutants using X-tremeGENE 9 and X-tremeGENE HP (Roche Applied Science, Penzburg, Bavaria, Germany) according to manufacturer’s protocol. Cells were plated on 25 mm glass coverslips coated with fibronectin bovine protein (Life Technologies, Carlsbad, CA) following Azioune et al. (20).

**Cell plating and surface patterning.** For plating, cells were dissociated using Accutase (Life Technologies) and plated on 25 mm glass coverslips coated with fibronectin bovine protein (Life Technologies, Carlsbad, CA) following Azioune et al. (20). 40 nm long crossbow fibronectin micropatterned cover slips were fabricated following protocol (*82*) using PLL-g-PEG purchased from (Surface solutions, Switzerlan), a UV lamps (UV ozone oven 185 nm equiped with ozone catalyzer, UVO cleaner, model 342-220, Jelight) and a chrome mask (Toppan).

**Single molecule Imaging.** All experiments were imaged with a Metamorph (Molecular Devices, Eugene, OR) controlled IX71 Olympus inverted microscope, a 100X objective with NA 1.45 (Olympus, Melville, NY) and an ILAS2 azimuthal TIRF FRAP head (ilas2; Roper Scientific, Tucson, AZ) in an azimuthal TIRF configuration. Cells were kept at 37°C in 5% CO2 with a heating chamber (Pecon, Meyer Instruments, Houston, TX). Single molecule movies of the red form of mEOS2 were imaged with a 561 nm laser (Cobolt Jive 150, Hubner) of incident power of 2kW/cm^2^, and a brightline quad-edge beamsplitter (Semrock Di01-R405/488/543/635). Photoconversion of mEOS2 was done with a 405nm laser (Stradus 405, Vortran) in a TIRF configuration. Imaging of iRFP was done with a 642nm laser (Stradus 642, Vortran) the same Brightline dichroic and a far red emission filter (BLP01-635R-25, semrock).

**Analysis of nanoclusters and trajectories**. We used the SLIMfast Matlab code (*83*) to recover single molecule localizations and DBSCAN to identify nanoclusters. Trajectories were reconstructed by finding the optimal global assignment between points in consecutive frames using an inference approach. The mapping of diffusivities in single cells was achieved by a Maximal Likelihood approach. Single cells map were averaged using custom built Matlab routines. All these procedures are detailed in the **supplementary information**

**Determination of membrane shuttling rates.** The shuttling rate of Rac1 to the membrane was analyzed by fluorescence recovery after photo bleaching of the whole basal membrane of the green form of mEOS2 in TIRF mode and recovery rates were determined as ca. 6 and ca. 20 min (**figure S1**) for spreading and spread cells, respectively.

**Rac1 FRET biosensors.** We established a stable cell line of 3T3 cells expressing a Rac1-FRET-biosensor (*49*). For imaging, cells were plated on glass coverslips with crossbow micropatterns. After 4 hours of adhesion, cells were imaged by epifluorescence using a Luca R camera (Andor on an Olympus IX71 microscope with a 60x magnification objective (Olympus PlanApo 60x NA 1.45). The same excitation and dichroic mirrors (ex: FF02-438/24, BS: FF-458-DiO2, Semrock) were used for the sequential acquisition of donor and acceptor images. A filterwheel was used to switch emission filters of donor (mCerulean, Em : FF01-483/32) and FRET acceptor (Em : FF01-542/27). Image processing included registration, flat-field correction, background subtraction, segmentation and FRET/donor ratio calculations. FRET ratio images were then aligned and averaged as described in the **supplementary information**.

**Optogenetics.** Recruitment of the catalytic domain of Tiam1 was performed via a Cry2-CIBN light gated dimerization as explained in reference (*45*). Localized recruitment was performed with 491 nm light that is highly effective for optogenetic recruitment but less efficient for photoconvertion of mEOS2. Recruitment laser pulses were applied every 10 seconds for ten minutes. Single molecule movies were obtained before and after recruitment. The low 405nm laser intensities used to photoconvert mEOS2 from the green to the red form did not introduce extensive global recruitment of Tiam1-Cry2-iRFP to the basal membrane. Imaging of iRFP was done with the same Brightline dichroic and a far red emission filter (BLP01-635R-25, semrock) and differential interference contrast (DIC) imaging was performed with a far red filter in the illumination path to avoid CRY2 recruitment.

**Inmunofluorescence.** Cell fixation and permeabilization was performed with 4% paraformaldehyde for 15 min and with 0.1% Triton X-100 or 0.5% NP40 for 5 min, respectively. To detect mouse WAVE2, a specific antibody named WP2 was raised against the peptide (C)NQRGSVLAGPKRTS in rabbits. Specific antibodies from the rabbit serum were affinity purified on a sulfolink column (Pierce) displaying the same peptide. WP2 recognizes murine WAVE2 by western blot, immunofluorescence and immunoprecipitates the WAVE complex. Anti-PIP3 was purchased from Echelon (Product number Z-P345b) and used in a 1:100 concentration for 60 minutes. Goat anti-mouse and goat anti-rabbit Alexa Fluor 647 labeled secondary antibodies were purchased from ThermoFisher (product numbers A-21236 and A-21245 respectively) and used in a 1:200 concentration for 60 minutes.

## Acknowledgments

We thank Leo Valon for designing and making the TIAM_linker_CRY2_IRFP optogenetic plasmid. We thank Fred Etoc for the initial mEOS2-Rac1 construction and for preliminary experiments. M.C. acknowledges financial support from French National Research Agency (LICOP n° ANR-12-JSV5-0002-01). M.C. and M.D. acknowledge funding from French National Research Agency (ANR) Paris-Science-Lettres Program (ANR-10-IDEX-0001-02 PSL), Labex CelTisPhyBio (N° ANR-10-LBX-0038) and France-BioImaging infrastructure supported by ANR Grant ANR-10-INSB-04 (Investments for the Future).

